# Functional Genomic Profiling of Schizophrenia-Associated Genes Reveals Key Microglial Regulators

**DOI:** 10.1101/2025.11.14.688490

**Authors:** Joy E. Horng, Liam T. McCrea, Rebecca E. Batorsky, Joshua J. Bowen, Camilla Boschian, Yoonjae Song, Roy H. Perlis, Steven D. Sheridan

## Abstract

Microglia are increasingly recognized as key regulators of neural circuit development and putative contributors to the pathophysiology of neuropsychiatric disorders such as schizophrenia (SCZ). However, the functional impact of SCZ-associated genes in microglia remains largely unexplored. Here, we performed an arrayed CRISPR targeting screen of 30 schizophrenia-associated genes predicted to be differentially expressed in human microglia-like cells. Target genes were prioritized based on post-mortem transcriptomic relevance and predicted ontology-based roles in phagocytosis pathways. We quantified phagocytic activity and morphological changes following gene targeting using high-content confocal imaging. Key targets, including *CYFIP1, MSR1, TREM2, SYK, ITGB2*, *ITGAM* and *IRF8*, modulated phagocytosis and altered morphological properties consistent with activation states, validating their functional roles in microglia. To elucidate transcriptional impact, we further applied a multiplexed RNA sequencing platform across gene targets. These analyses revealed gene-specific transcriptional signatures, implicating divergent pathways related to phagocytic, activation, cytoskeletal, and lysosomal function. Together, these findings demonstrate the utility of CRISPR-based functional genomics in characterizing microglia function and identifying new target genes and mechanisms that may underlie their contributions to schizophrenia pathophysiology.

## Introduction

Schizophrenia is a genetically complex, highly heritable neuropsychiatric disorder characterized by symptoms including psychosis, cognitive deficits, and social withdrawal, typically emerging in late adolescence or early adulthood.(1, 2) Schizophrenia is also highly polygenic(3): Genome-wide association studies (GWAS) have identified hundreds of schizophrenia-associated loci, including several within immune-related genes (4–6), suggesting an overlap between synaptic and neuroimmune pathways.(4, 5) Convergent evidence for neuroimmune mechanisms comes from large-scale postmortem RNA sequencing studies(7), suggesting some risk genes are highly expressed in microglia.(8, 9)

These brain-resident innate immune cells are key regulators of neural development and function.(10, 11) They mediate synaptic pruning via complement-dependent and less well-understood mechanisms to maintain homeostasis through surveillance and phagocytosis.(12–15) Postmortem, live imaging, and transcriptomic studies implicate increased inflammatory gene expression, and altered microglial morphology and activity in schizophrenia.(16–20) Further, patient-derived *in vitro* models demonstrate excessive microglial synaptic engulfment in microglia-like cells derived from individuals with schizophrenia (21), yet the functional consequences of disrupting schizophrenia-associated genes in human microglia remain unknown.

To facilitate the investigation of human microglia, multiple *in vitro* cellular models have been developed using either induced pluripotent stem cell (iPSC) differentiation (22–24) or direct transdifferentiation from peripheral blood mononuclear cells (PBMCs) (21, 25, 26), which permit higher throughput screening (27) and investigation of disease-relevant phenotypes. CRISPR-based platforms, such as pooled screens of the druggable genome in Cas9-expressing engineered iPSC-derived microglia-like cells, have recently identified genes involved in survival and function.(28) However, despite extensive CRISPR-based functional genomics investigation in neurons (29–31), systematic *in vitro* genetic screens targeting schizophrenia-risk genes in microglia remain scarce.

In this study, we leveraged an arrayed CRISPR targeting approach in PBMC-derived human microglia-like cells (piMGLCs) to investigate the functional roles of schizophrenia-associated genes. The list of target genes was identified based on postmortem differentially expressed genes (DEGs) and expression in human microglia, with further prioritization using predicted involvement in phagocytosis-related pathways from Gene Ontology enrichment analyses. We first conducted primary functional screening, which assessed the capacity of piMGLCs to phagocytose isolated human synaptosomes in an immunocytochemistry and high-content confocal imaging-based *in vitro* model of synaptic pruning.(21, 25–27) Selected genes that altered phagocytosis upon gene targeting were then selected for orthogonal secondary screening using redesigned sgRNAs to validate their effects on phagocytic activity and measure indicators of activation state, including cellular morphology and marker expression. Finally, we employed a high-throughput RNA sequencing platform (32) to profile the transcriptional impact across gene targets, enabling the discovery of downstream pathways and potential mechanisms of action. We hypothesized that specific schizophrenia-associated genes would modulate microglial activity, thereby contributing to the synaptic alterations observed in disease in postmortem and *in vitro* studies. Beyond offering novel insights into microglia-mediated mechanisms in schizophrenia, this approach establishes a scalable platform for further higher throughput neuroimmune genetic screening.

## Methods and Materials

### PBMC-Derived Induced Microglia-Like Cell (piMGLC) Culture and Seeding

piMGLCs were derived as previously described (27) with some modifications. Briefly, frozen PBMCs (Vitrologic, Inc. for primary screen, Charles River, Lowell MA; Hemacare, Northridge CA for secondary screen) were thawed at 37°C and transferred into RPMI-1640 (Sigma, #R8758) supplemented with 10% heat-inactivated FBS (Sigma, #12306C) and 1% Penicillin/Streptomycin (Life Technologies, #15140-122). After centrifugation at 300 g for 5 min (brake off), cells were resuspended, counted, and plated at ∼500,000 cells/cm² in 6-well tissue culture plates (Corning, #353046) pre-coated with Geltrex (Gibco, #A1413202) for 1 hour. After 24 hours, media was replaced with RPMI-1640 containing 1% Penicillin/Streptomycin, 1% Glutamax (Life Technologies, #35050-061), 100 ng/ml IL-34 (Biolegend, #577904), and 10 ng/ml GM-CSF (PeproTech, #300-03). Cells were incubated for 8 days to induce trans-differentiation.

### Ribonucleotide protein (RNP) delivery of gRNAs

Guide RNAs (gRNAs) (Synthego library; Table S2) were resuspended to 25 μM. SpCas9 protein (IDT, #1081059) was diluted to 31 μM in Duplex buffer (IDT, #11-01-03-01) and combined with gRNAs at a 1:1.5 molar ratio (SpCas9:gRNA) to form ribonucleoprotein (RNP) complexes (∼10.8 μM final concentration). Complexes were gently mixed by pipetting, incubated for 10 min at room temperature, diluted to 7.2 µM, and kept on ice until use as 5μl aliquots in 8-strip PCR tubes per 25,000 cells (per reaction).

piMGLC culture supernatant was collected, filtered through a 0.22 µm Steriflip-GP unit (EMD Millipore, #SCGP00525), and retained as Conditioned Medium for post-nucleofection culture. piMGLCs were washed with DPBS and detached using Accutase (Sigma, #A6964) for 5 min at 37 °C, centrifuged (300 g, 5 min), and resuspended in fresh medium.

For nucleofection, 25,000 cells (for assays) or 75,000 cells for Drug-seq were resuspended in 20 µl P3 buffer (Lonza P3 Nucleofection Kit, #V4SP-3096) and mixed with prepared RNPs. Cell-RNP suspensions were transferred into nucleofection cuvettes and nucleofected (Lonza 4-D nucleofector) using program EA-100. After a 10 min recovery at room temperature, cells were seeded into Geltrex-coated 96-well plates (Corning, #3904) with 180 µl Conditioned Medium (total 200 µl per well) and cultured for 6 days.

### Phagocytosis assays

Phagocytosis assays were conducted in 96-well plates (Corning, #3904) seeded with piMGLCs at a density of 60,000 cells/cm² (20,000 cells per well in 200 µl). Human synaptosomes were thawed at room temperature and mixed 1:1 with 0.1 M sodium bicarbonate (pH 9), then labeled with pHrodo-Red dye (Invitrogen, #P36600, 6.67 µg/µl) at a protein:dye ratio of 2:1 for 1 hour. Labeled synaptosomes were washed by adding PBS (pH 7.4), pelleted at 12,000 rpm for 15 minutes, and resuspended in basal RPMI-1640 at 0.15 mg/ml. Synaptosomes were then sonicated in a Branson 1800 (Emerson, #M1800) at 40 kHz for 1 hour. Synaptosomes were added to each well at 3 µg/well using a multichannel pipette. After 3 hours, assays were terminated by fixation with 4% paraformaldehyde (Electron Microscopy Sciences, #15713S).

### RNA-seq data analysis

Raw paired-end FASTQ files were trimmed with Trim Galore v0.6.10 (33) and aligned to hg38 using STAR v2.7.11b.(34) The demultiplexed count matrix contained an average of 1.51 ± 0.39 million reads per sample. Differential expression was analyzed with DESeq2.(35) For CRISPR-targeted genes, significance was defined as unadjusted p < 0.1, while for all other genes, FDR-adjusted padj < 0.1 was used, based on the rationale that expression of the perturbed gene is expected to change, reducing the likelihood of false positives relative to transcriptome-wide tests. Functional enrichment of DE genes with >10 DEGs per target was performed using clusterProfiler enrichGO.(36) Code is available at https://github.com/mgb-cqh/crispr30.

Additional methods and details available in Supplemental Material and Methods.

## Results

### Targeted functional genetic screening in human primary PBMC-derived reprogrammed microglia-like cells identifies genes that modulate synaptosome phagocytosis

To curate schizophrenia liability genes, we focused on genes which are both expressed in microglia (9, 37, 38) and found to be differentially expressed in schizophrenia in postmortem studies (7, 8, 39–41), particularly in the dorsolateral prefrontal cortex (DLPFC) (42). We curated a set of ∼200 genes from these datasets (Table S1) and selected those determined by gene ontology analysis (GO, KEGG, GSEA) to be involved in phagocytosis, resulting in 30 genes prioritized for initial screening. (Table S1).

Due to the primary nature of the PBMCs, which cannot be passaged like cell lines engineered to express Cas9 protein (43–45), we adapted a modified protocol (46, 47) for CRISPR-mediated genome editing using Cas9/sgRNA ribonucleoprotein (RNPs) delivery of purified recombinant Cas9 complexed with sgRNAs, resulting in efficient manipulation of viable PBMC-induced microglial-like cells (piMGLCs) for use in screening of schizophrenia-related genes.

Using this arrayed RNP-based CRISPR system with three sgRNAs per target gene (Table S2) per well in 96-well plate format, we performed gene targeting in piMGLCs (Fig. S1). 6 days post sgRNA delivery, we performed phagocytosis screens (Fig. 1a) using pHrodo dye-labeled human iPSC-derived neuronal synaptosomes in a model of synaptic pruning (21, 26), followed by immunocytochemistry (ICC) to identify microglia by IBA1+, and high-content confocal imaging to quantify phagocytic index (see Methods). This primary screen revealed seven potential repressors of phagocytosis (i.e. gene targeting *increased* phagocytosis) compared to the non-targeting negative control sgRNAs (NEG) (Fig. 1b, Table S3). Conversely, the majority of genes reduced phagocytic activity upon gene targeting.

**Figure 1.**
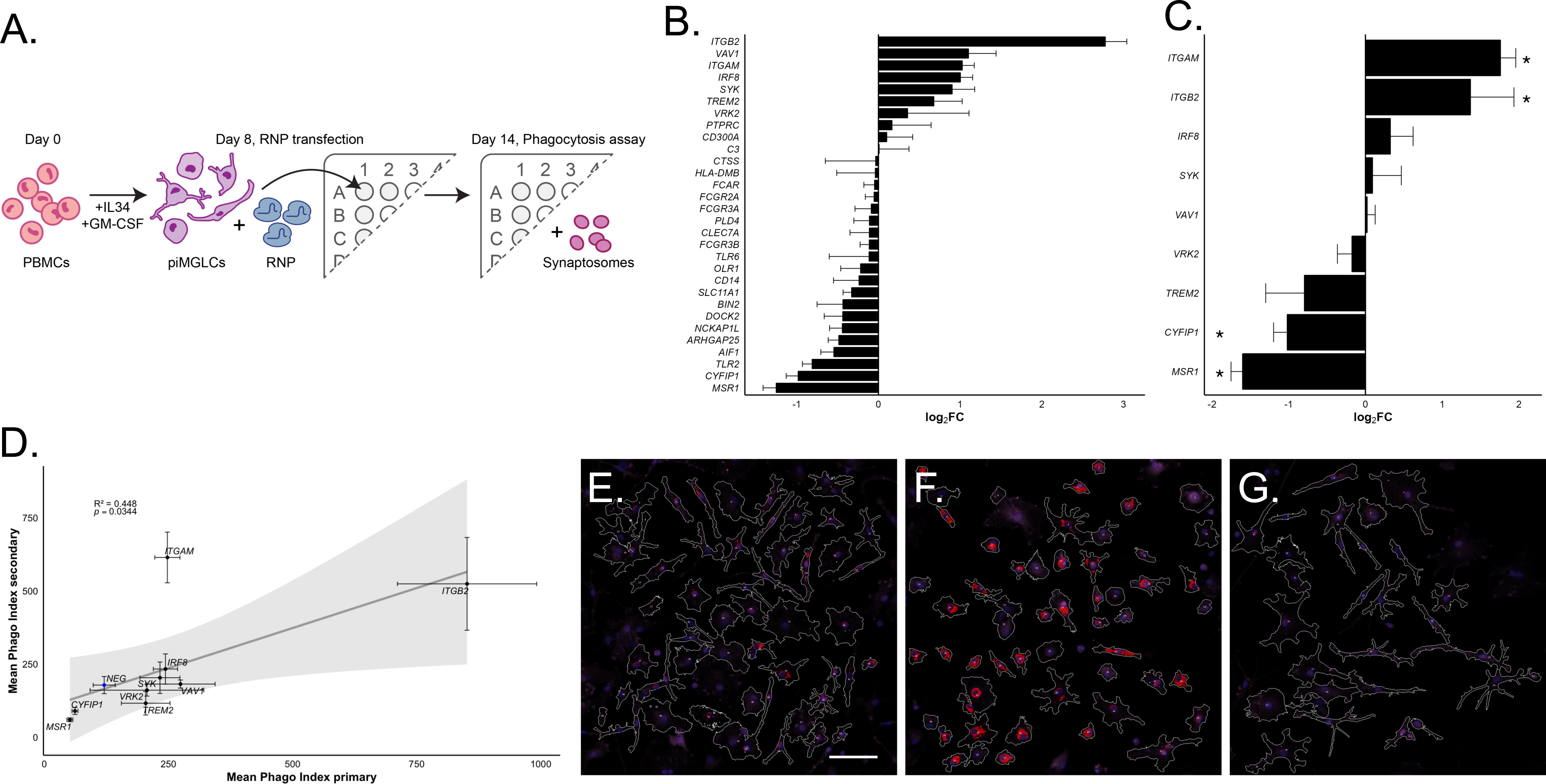
Image-based arrayed CRISPR screen for synaptosome phagocytosis modulators in piMGLCs. (a) Overview workflow for derivation of piMGLCs, ribonucleoprotein (RNP) transfection and assay. (b) Phagocytosis primary screen of 30 curated schizophrenia associated genes. Phagocytic index indicated as fold-change (FC) compared to non-targeting gRNA negative control. (*n*=3 replicate well mean cellular values, # cells per well in Table S3, error bars indicate SEM). (c) Phagocytosis secondary screen of 9 schizophrenia associated genes selected from primary screen. Phagocytic index indicated as fold-change (FC) compared to non-targeting gRNA negative controls. (*n*=3 replicate well mean cellular values, # cells per well in Table S4). (d) Correlation of phagocytic indices between primary and secondary screens. Error bars indicate SEM, grey area indicates 95% confidence. (e-g) Representative masked cell images used for quantification, white mask outline containing pHrodo stained synaptosomes (red) corresponding to (e) non-targeting gRNA negative control and gRNAs targeting (f) *ITGB2* and (g) *MSR1*. Scale bar = 100μM

To determine independent reproducibility and confirm findings from the primary screen, secondary screening was conducted on selected target genes found to alter phagocytic function in the primary screen using redesigned, orthogonal sgRNAs (two per gene, Table S2) and additional negative controls, including an eGFP sgRNA (eGFP gene is not present in these cells) added to the non-targeting gRNA sequence (NTS) used in the primary screen. Consistent with primary results, *ITGB2* and *ITGAM* targeting increased phagocytosis while *MSR1* and *CYFIP1* targeting reduced phagocytosis (Fig. 1c, Table S4).

Correlation between primary and secondary screening results revealed a correlation for most gene sgRNA treatments (R^2^=0.448, across genes, Fig. 1d) other than *ITGAM.* Excluding *ITGAM,* which by visual inspection was a marked outlier, primary and secondary results were highly correlated (R^2^=0.936 excluding *ITGAM*, Fig. S2), possibly reflecting a difference in editing efficiency between the sgRNAs used in the different screens.

### Phenotypic screening identifies genes that modulate microglial morphology and activation state

We previously demonstrated that the morphometric parameters of solidity and eccentricity can be used to indicate distinct morphological classes of microglia: ramified, amoeboid, and intermediate bipolar/rod-shaped.(27, 48) Microglia typically reside in a surveillant state in the adult brain, characterized by a ramified morphology, (49) but they alter their morphology upon a change in functional state. This allows morphology to serve as a proxy for activation status, with ramified cells representing a surveillant state and amoeboid cells indicating one of the multiple known microglial activation states.(50)

To assess these morphological transitions due to select gene targeting, we used our arrayed image-based screening platform to simultaneously measure morphometric parameters and phagocytic function in both the primary and secondary screens (Fig. 2). We analyzed these piMGLCs using CellProfiler (51), extracting solidity and eccentricity values to infer microglial activation states by morphology at scale.

**Figure 2.**
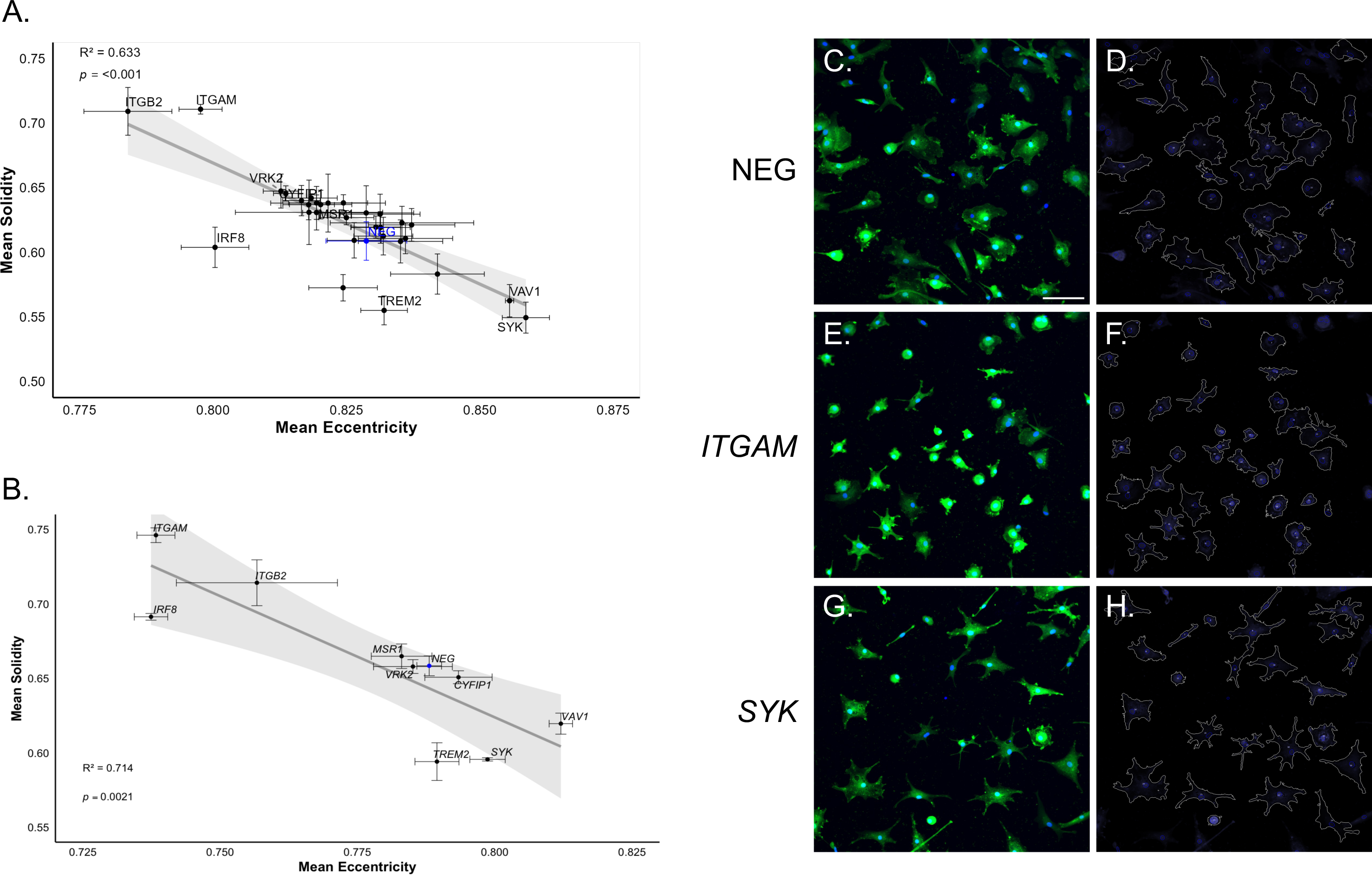
Phenotypic screening identifies genes that modulate piMGLC morphology. (a) Morphometric parameters solidity versus eccentricity for 30 indicated CRISPR targeted genes in primary screen. (*n*=3 replicate well mean cellular values, # cells per well in Table S3). (b) Morphometric parameters solidity versus eccentricity for 9 CRISPR targeted genes selected from primary screen. (*n*=3 replicate well mean cellular values, # cells per well in Table S4). Error bars indicate SEM, grey area indicates 95% confidence. (c,e,g) Representative images (IBA1+ green, nuclei blue) and (d,f,h) corresponding masked cell outlines for quantification of indicated non-targeting gRNA negative control (NEG) and gRNAs targeting *ITGAM* and *SYK*. Scale bar = 100μM.

In the primary screen, we observed pronounced changes in cellular morphometric parameters following targeting of several genes, including *ITGB2, ITGAM, SYK, VAV1, TREM2, IRF8,* and *VRK2* (Fig. 2a, Table S3). Targeting these genes prompted distinct morphological changes exhibiting shifts towards a ramified (high eccentricity, low solidity) or amoeboid (low eccentricity, high solidity) morphology. For instance, targeting of *ITGB2* and *ITGAM*, both components of the complement receptor 3 (CR3) (52), each exhibited increased solidity and reduced eccentricity relative to negative controls, consistent with a more amoeboid morphology. Other gene targets such as *VAV1* and *SYK* resulted in more ramified morphology, demonstrating varied effects on morphology across gene targets.

Further morphometric analyses of the secondary screen indicate the morphological alterations observed in the primary screen were robustly reproduced (Fig. 2b-h, Table S4). We observed strong correlations between the two screening rounds for both solidity and eccentricity (R² = 0.82 and R^2^ = 0.708, respectively, Fig. S3), indicating reproducible morphology phenotypes upon gene-specific disruption across biological replicate screens.

We further found that morphology correlated with phagocytic index (Figs. 3a-b). Specifically, increased eccentricity was associated with lower phagocytosis (R^2^ = 0.479), while increased solidity generally associated with higher phagocytosis (R^2^ = 0.564) across the genotypes examined. Overall, more amoeboid, less ramified microglia tended to be more phagocytic.

**Figure 3.**
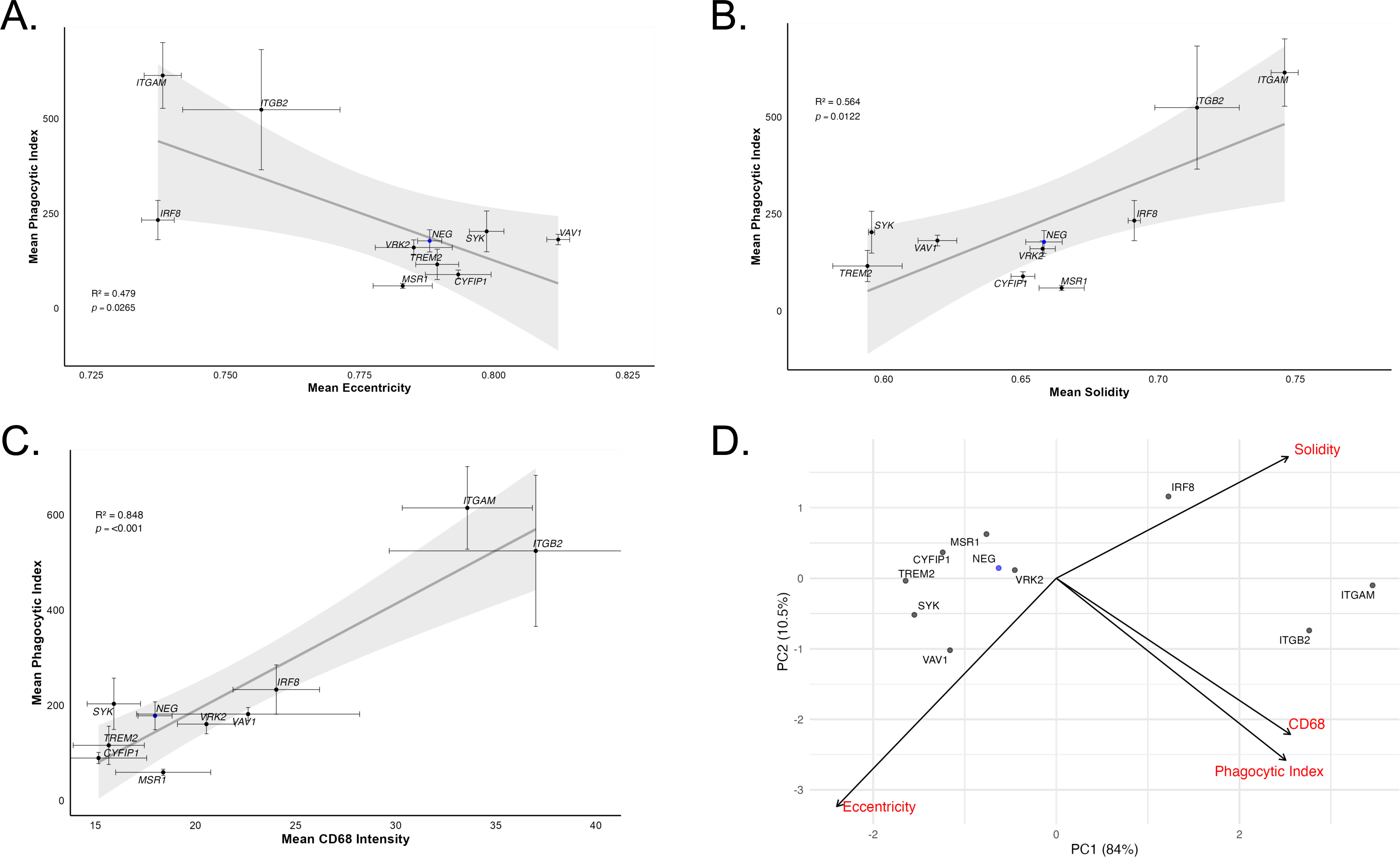
Functional and phenotypic analyses between targeted genotypes indicate correlations between phagocytic activity, morphology, and microglial activation marker expression. (a) Increased eccentricity results in lower phagocytosis while (b) increased solidity correlates with higher phagocytosis. (c) Correlation between phagocytic index and CD68 expression. (*n*=3 replicate well mean cellular values, # cells per well in Table S4). Error bars indicate SEM, grey area indicates 95% confidence. (d) Multi-dimensionality reduction using principal component analysis (PCA) of solidity, eccentricity, phagocytosis, and CD68 marker expression enables discrimination between gene targeted groups.

To further explore relationships between targeted genotypes and functional states, we incorporated the expression level of the activation marker CD68 (53, 54) into our evaluations of microglial activation. CD68 expression levels were highly dispersed across genotypes (Fig. 3c), indicating specific gene targets lead to increased (e.g. *IRF8, ITGAM* and *ITGB2*) or decreased (e.g. *TREM2, CYFIP1* and *SYK*) activation compared to negative controls. Furthermore, CD68 expression positively correlated (R^2^ = 0.848) with phagocytic activity, suggesting coordinated changes between activation and phagocytic function.

We applied dimensionality reduction (Fig. 3d) using principal component analysis (PCA) of solidity, eccentricity, phagocytosis, and CD68 expression to understand the relationship among these phenotypes. These multidimensional phenotypic profiles separate functional microglia states that result from targeting genes. For example, *ITGAM* and *ITGB2* targets clustered upon PCA, exhibiting high activation and phagocytic capacity profiles, while *CYFIP1*, *TREM2* and *MSR1* demonstrate high similarity due to lower activation and phagocytic function. This approach highlights that measures of microglia state cannot be captured by individual parameters. It further emphasizes the potential for high-throughput image-based assays to reveal unexpected or nuanced gene function relevant for prioritizing further disease modeling efforts such as clonal gene knockout in human iPSCs-derived microglia.

### Transcriptomic Profiling Reveals Diverse Gene Targeting Effects on Microglial State

We next performed DRUG-seq (32, 55), a multiplexed RNA-sequencing approach optimized for high-throughput screening, to investigate the transcriptomic consequences of individual gene targeting in piMGLCs.

For each of the 30 targeted genes, we generated three biological replicate libraries and compared their expression profiles with non-targeting negative control (NTCs) samples processed in parallel. Three of the targeted genes (*PLD4*, *FCGR3B,* and *FCAR*) were minimally expressed in the piMGLCs (average coverage in controls < 2 CPM, Fig. S4), and were thus removed from further analyses. Of the remaining expressed genes, six did not result in significant changes in their own transcript levels: *ARHGAP25*, *SLC11A1*, *C3*, *TLR6*, *OLR1* and *VRK2* (raw p-value <0.1, as shown by the diagonal in Figure 4a). This may indicate the selected gRNAs failed to cause nonsense-mediated decay (NMD) (56, 57) of their target transcripts, but may still alter protein function through out-of-frame indel induced truncations. The remaining targets (21/30) resulted in significant alterations in their respective targeted gene transcript levels compared to NTCs, with genes such as *ITGAM*, *NCKAP1L*, *BIN2*, *HLA-DMB* and *DOCK2* among the most reduced.

**Figure 4.**
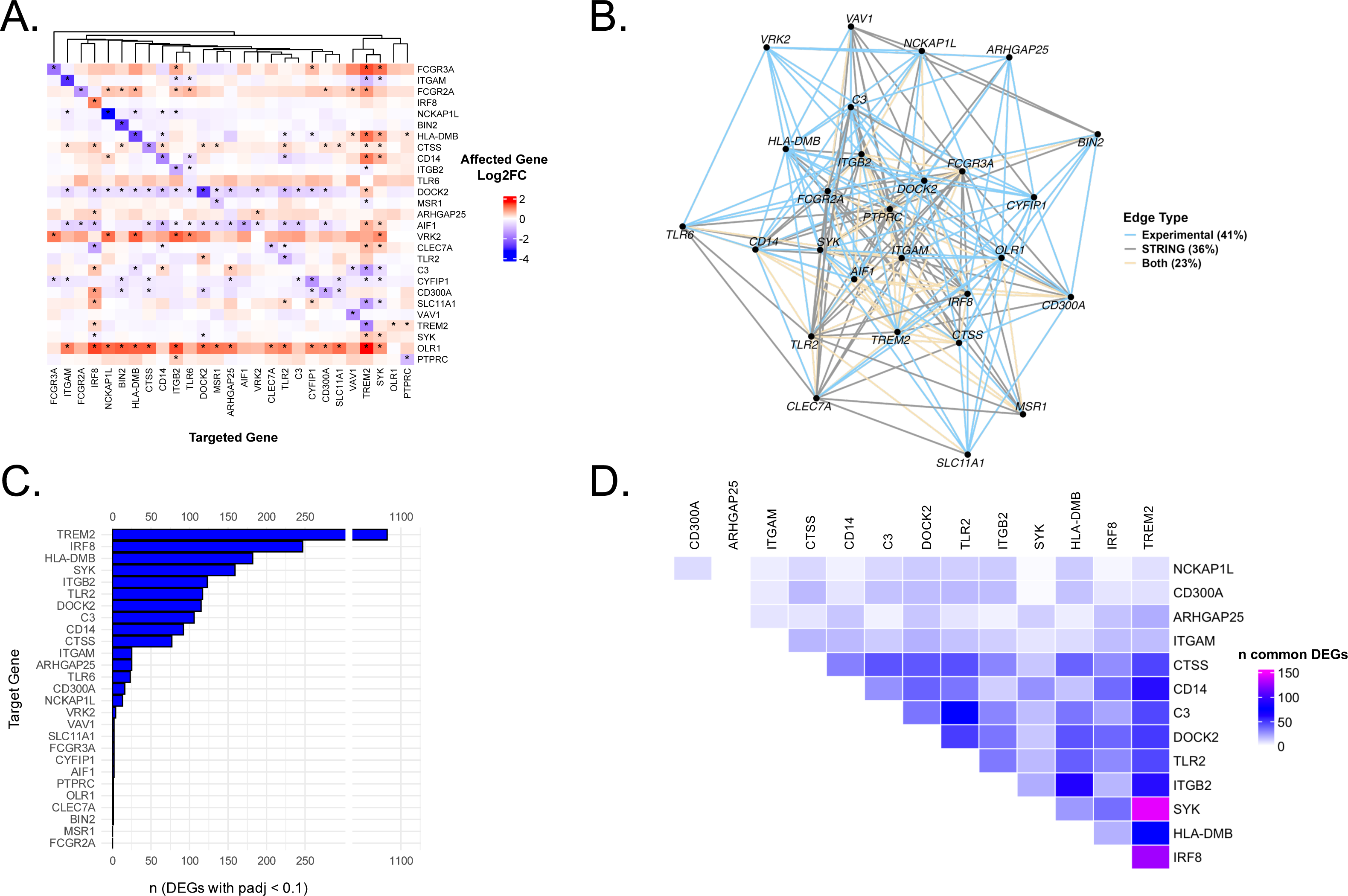
Differential Expression in Drug-Seq. (a) Pairwise effects of each target gene on expression of all other targeted genes. Colors show log2 fold change. Asterisks mark combination with p-value < 0.1. (b) Comparison with documented STRING-db pairwise gene interactions. Lines indicate (blue) current experimental only (41%), (grey) STRING-db only (36%), (yellow) both experimental and STRING-db (23%). (c) Number of DEGs (padj < 0.1) per target gene. Bar colors reflect the DEG status for that target gene (red = pvalue < 0.1 & log2FC > 0; blue = pvalue < 0.1 & log2FC < 0; black = otherwise). (d) Overlap of DEG in common between gene targets (minimum 5 DEG in common with at least one other gene target).

Interestingly, initial RNA sequencing indicated that targeting *IRF8* and *SYK* led to increased expression of their transcripts. However, further confirmation by qRT-PCR (Fig. S5) only supported this observation for *IRF8*, not *SYK*. Similarly, Western analysis of these gene targets indicated that IRF8 protein level was slightly increased, while protein expression of SYK was markedly reduced. These results suggests possible alternative translation initiation (ATI) leading to increased transcript stability (57) in the former, and/or lack of nonsense mediated transcript decay (NMD) in the latter (56, 57). While this does not rule out the introduction of non-functional mutations in either of these proteins, it remains likely that both proteins were impacted given our pronounced functional and phenotypic observations.

To examine potential regulatory relationships among the targeted genes, we cross-referenced expression of all 30 target genes across the DEG results from each perturbation, generating a pairwise differential expression matrix (Fig. 4a). The targeting of several genes (e.g. *TREM2*, *SYK* and *IRF8*) triggered broad transcriptomic changes in other targets, indicated by significant log fold changes (FC). Alternatively, expression differences in some affected genes (e.g. *DOCK2*, *AIF1* and *OLR1*) were induced by many targeted genes, suggesting downstream regulation by other targeted genes. Many of these pairwise comparisons (23%) corresponded to previously documented interactions in the STRING database (58) (Fig. 4b). In contrast, 41% of the identified relationships have not been reported in STRING and may represent microglia-specific regulatory networks.

Further transcriptomic analysis identified genes that exert the most profound impacts on the transcriptome following CRISPR targeting. Several gene targets are notable for driving the largest numbers of significantly differentially expressed genes (DEGs) (Fig. 4c, Table S6), such as *TREM2*, *IRF8*, *HLA-DMB* and *SYK*, representing loci whose perturbation results in widespread transcriptional reprogramming. Notably, targeting *ITGB2* resulted in approximately 5-fold more DEGs than *ITGAM*, despite both encoding subunits of the CR3 receptor (52), suggesting non-overlapping functions.

Pairwise analyses that show the most significant overlap in DEG profiles (Fig. 4d) highlight potential candidate interactions or shared pathways. The most notable overlaps, indicated by the largest intersecting DEGs (e.g., *TREM2* and *SYK* are genes within the same phagocytic and activation pathways (59)) emphasize gene pairs whose parallel activity may contribute to pathway outputs or whose combined loss uncovers new regulatory dependencies.

The biological implications of these findings suggest gene targets that yield the highest number of DEGs may serve as regulatory hubs or critical nodes within microglial functional pathways, and that gene pairs with maximum DEGs overlap identify possible points of redundancy in gene networks. Identifying such genes and pairs informs prioritization for mechanistic follow-up, functional assays, or therapeutic targeting.

### Functional Programs Induced by Targeted Genes

GO Biological Process enrichment highlighted functional themes relevant to microglial biology. We curated the enriched processes into a set of broad functional categories that are biologically relevant and represented across targets, including chemokine/cytokine signaling, lysosomal pathways, phagocytosis, cell death, cell adhesion, actin cytoskeletal remodeling, metabolism, protein production, complement activation, and autophagy (Fig. 5a, Table S7).

**Figure 5.**
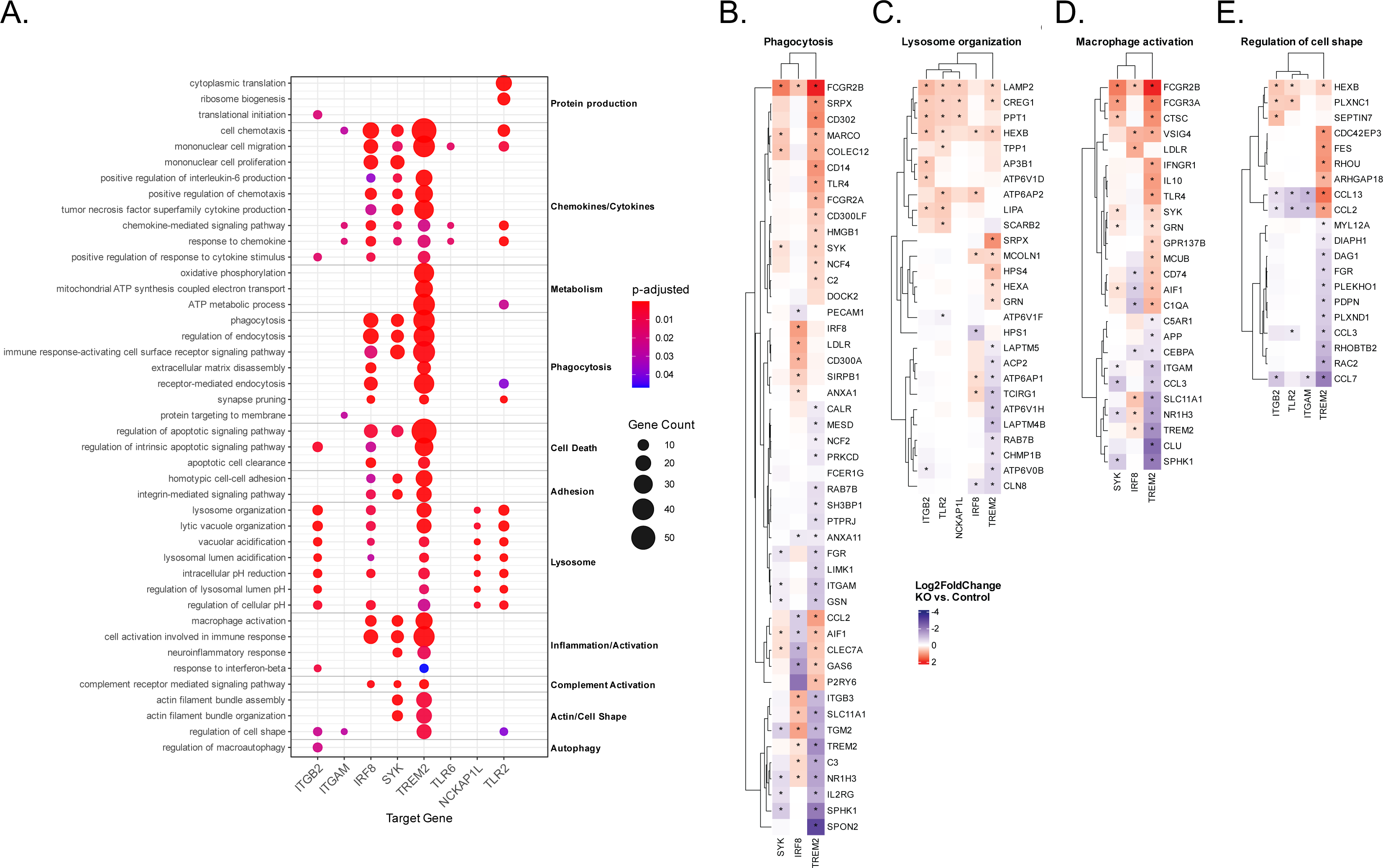
Transcriptomic analysis of targeted genes (a) Select GO Biological Process enrichment for target gene DEG. Dot color encodes adjusted p-value. Dot size encodes term gene count. (b-e) Gene-level differential expression heatmaps for select GO Biological Process terms enriched in the DEGs of target genes (b) phagocytosis, (c) lysosome organization, (d) macrophage activation and (e) regulation of cell shape. Log2FC = log fold change (Targets vs Control); red indicates up-regulation and blue indicates down-regulation. Asterisks mark genes with |log FC| > 0.2 and adjusted *p*-value (padj) < 0.1. Only treatments/genes with available data are displayed.

Targeting *TREM2* and *SYK* affected processes central to microglial function including phagocytosis, lysosome organization and acidification, cell activation, cytokine/chemokine signaling, and actin cytoskeletal remodeling (59–61), supported by our functional and phenotypic data. *IRF8* and *TREM2* targeting affected the expression of numerous other targeted genes (Fig. 4d) and a broad set of GO processes, confirming central roles in microglial regulation. (62) *ITGB2* and *ITGAM*, despite encoding subunits of the CR3 receptor (CD11b and CD18, respectively) (14, 63), showed divergent enrichment patterns due to *ITGB2*’s broader transcriptomic impact (Fig. 4c).

Beyond these major regulators, targeting *TLR2* impacted protein production and lysosomal acidification processes, indicating effects on fundamental cellular processes such as translation and degradation. *NCKAP1L* targeting showed more specialized enrichment, particularly in lysosomal acidification and actin cytoskeletal organization, consistent with its known role in the WAVE complex and phagocytosis.(64, 65)

We further compared gene targets by examining the gene-level changes in four strongly enriched processes (Figs. 5b-e). First, phagocytosis was a strongly enriched process across *SYK*, *IRF8,* and *TREM2* targets (Fig. 5a). *TREM2* affected the largest number of genes (Fig. 5b) in this category (39), including upregulation of canonical phagocytic receptors and effectors such as *FCGR2A/B*, *CD14*, *MARCO*, *CD302*, *CLEC7A*, as well as downregulation of *ITGAM* and *C3*. *IRF8* targeting influenced a partially overlapping set of 18 genes, but with predominantly opposite directionality, aside from consistent upregulation of *FCGR2B* and consistent downregulation of *ANXA11*. Targeting *SYK* affected 13 phagocytosis genes, including *FGR*, *MARCO*, *CLEC7A*, and *ITGAM*, and the direction of expression changes were broadly concordant with those observed in *TREM2*.

Lysosomal organization was also affected across multiple targets, including *ITGB2*, *TLR2*, *NCKAP1L*, and *TREM2*, highlighting disruption of phagolysosomal processes (Fig. 5c). Several lysosomal genes, including *LAMP2*, *CREG1*, and *HEXB* were consistently upregulated across multiple targets, potentially reflecting lysosomal stress. In contrast, a broader collapse of lysosomal organization is represented by the following: *ATP6V0B* was downregulated in *ITGB2* and *TREM2* targets which indicates reduced acidification (66), *CLN8* was reduced in *TREM2* and *IRF8* targets, suggesting deficits in trafficking, (67) and targeting *TREM2* exhibited the most extensive downregulation of lysosomal genes, including *CHMP1B*, *ACP2*, and multiple *LAPTM* family members.

Macrophage activation pathways were also prominently enriched across *TREM2, IRF8*, and *SYK* (Fig. 5d), with mixed overall effects on activation. Changes were generally modest upon targeting *IRF8*, with classical activation markers such as *C1QA*, *AIF1*, and *CD74* reduced, while homeostatic or anti-inflammatory genes including *VSIG4* and *LDLR* were increased. Targeting *SYK* and *TREM2* resulted in highly concordant transcriptional profiles and included strong upregulation of Fc receptor and antigen-processing genes such as *FCGR2B*, *FCGR3A*, *CTSC*, and *CD74*, as well as downregulation of inflammatory and metabolic regulators including *SPHK1*, *NR1H3*, *CCL3*, and *CLU*. This pattern suggests that *SYK* and *TREM2* disruption elicits a shared transcriptional program characterized by reduced proinflammatory activation and enhanced antigen-presenting functions, despite decreased phagocytic activity in functional assays.

Finally, regulation of cell shape was enriched across several targets (Fig. 5e), reflecting both signaling-mediated and cytoskeletal remodeling processes. *TREM2* had the broadest impact, altering both upstream signaling components and downstream effectors, including small GTPase regulators (*RHOU, RAC2, CDC42EP3, ARHGAP18*) and cytoskeletal modulators (*DIAPH1, MYL12A*). In contrast, *ITGB2, ITGAM*, and *TLR2* primarily influenced chemokine expression (e.g. *CCL13*, *CCL2* and *CCL7*).

## Discussion

In this study, we systematically investigated the functional roles of schizophrenia (SCZ)-associated genes in a human microglia-like cellular model using an arrayed CRISPR targeting screen coupled with high-content imaging and transcriptomic profiling. Our results identify a subset of microglial-expressed genes that regulate phagocytosis, cellular morphology, and activation states. These findings advance our understanding of microglial involvement in schizophrenia and demonstrate the utility of using integrated functional genomics to determine disease-relevant cellular mechanisms.

The genes prioritized for CRISPR screening were curated from SCZ postmortem transcriptomic datasets (7) and further enriched by microglial expression (8, 9) and phagocytic genetic ontogeny (DAVID (68, 69) and GSEA (70, 71)). This approach enabled the functional validation of these selected candidates within a human microglial context, connecting large-scale association studies with mechanistic insight.

To address the limitations in arrayed CRISPR screening that can introduce confounding factors, including variable well-to-well editing efficiencies from RNP nucleofection, potential variable expression of target genes, and potential off-target effects, we designed orthogonal sgRNAs for secondary validation of selected targets in addition to incorporating additional negative controls (e.g. non-targeting gRNAs and eGFP, the latter not present in these cells) to compare to for well-level image data normalization. This method ultimately presents a relevant, efficient primary cell screening method that can be used to identify and characterize the functional outcome of gene disruption to design subsequent genetic studies using clonal iPSC-derived microglia and/or 3D culture systems for greater fidelity and resolution of cell-autonomous versus intercellular effects.

Among the most robust impacts on phagocytosis were increased activity following targeting of *ITGAM* and *ITGB2* and reduced activity in *MSR1* and *CYFIP1*. Many genes that demonstrated impacts on phagocytic uptake of synaptosomes are components of known pathways implicated in synaptic pruning, a process hypothesized to be dysregulated in SCZ.(72–75) The altered phagocytosis observed upon their targeting supports this model and suggests that deficits in microglial engulfment may contribute to the synaptic dysfunction observed in patients.

Our application of high-throughput multiplexed transcriptomic analyses provided a transcriptome-wide view of CRISPR targeting-induced changes. It revealed substantial heterogeneity among genes, both in the magnitude of transcriptional impact and in the pathways affected. Building on the analysis of interacting gene pairs, these pathway-level data reveal how individual targets give rise to convergent transcriptional programs. While some targets, such as *TREM2*, *SYK*, and *IRF8*, elicited broad transcriptional responses affecting multiple cellular programs, including phagocytosis, chemokine signaling, and lysosomal processes, other genes, such as *ITGB2*, *ITGAM*, *NCKAP1L*, and *TLR2*, showed more focused process enrichment, often linked to core effector functions such as actin remodeling, lysosomal acidification, or protein production.

Phenotypic effects did not consistently parallel transcriptomic breadth, as some genes produced broad transcriptional changes but modest phenotypic shifts, such as *IRF8* and *TREM2*, while others showed strong functional responses and relatively focused transcriptomic signatures, such as *ITGB2* and *ITGAM. SYK* uniquely exhibited strong transcriptional and phenotypic effects, consistent with its role as a central signaling hub. This discordance highlights that different genes can influence microglial function through distinct regulatory modes, some acting via specific effector pathways that directly impact cellular behavior, and others through broader transcriptional networks that modulate microglial states in more complex ways. These insights point to distinct regulatory circuits disrupted by each gene and suggest potential points of convergence or synergy.

The integration of phenotypic and transcriptomic datasets supports a model in which subsets of schizophrenia-associated genes control microglial function through distinct mechanisms, some reducing phagocytic capacity directly through receptor or lysosome function, (e.g., *ITGB2*, *IRF8*, *TREM2*, *NCKAP1L* and *TLR2*), others also altering cellular activation or survival pathways (e.g., *IRF8*, *TREM2* and *SYK*) and/or through direct transcriptional regulation via transcription factors (e.g., *IRF8*). For instance, targeting *IRF8* led to increased expression of pro-inflammatory transcripts, suggesting an activation response rather than a direct phagocytic defect, thus highlighting the importance of multiplexed phenotyping strategies when interpreting the impact of gene perturbations in complex cellular systems. Other genes in this study, such as *VRK2*, that resulted in functional and phenotypic disruptions upon targeting with less understood connections to these other schizophrenia-implicated genes, warrant further investigation.

Our results support a model in which schizophrenia-associated genes modulate key microglial functions, including phagocytosis and immune activation, through distinct molecular mechanisms. More broadly, these findings present a scalable platform for functional genomics in microglia, enabling disease modeling and therapeutic discovery applicable to schizophrenia, and neurodevelopmental or neurodegenerative disorders. The identification of regulatory hubs and redundant gene networks provides key targets for further mechanistic follow-up and potential intervention strategies.

## Author Contributions

J.E.H. and L.T.M, optimized the batch differentiation and characterization of piMGLCs, and scaling phagocytosis assay to 96-well format. J.E.H. developed and optimized the large-scale RNP CRISPR procedure for gene targeting in piMGLCs and performed the phagocytosis screens and phenotypic assays. L.T.M. generated, purified, and characterized large scale synaptosome preparations. L.T.M. and J.E.H. developed CellProfiler pipelines and performed data analysis. R.E.B. performed transcriptomic analyses and interpretation of data. J.J.B and Y.S. performed gene expression and Western analyses. J.J.B. and C.B. performed sample and multiplexed Drug-seq library preparation for sequencing. S.D.S., R.E.B., and R.H.P wrote the manuscript. S.D.S and R.H.P conceived and directed the project. All authors have read, provided edits, and approved the manuscript for publication.

## Funding

This work was supported by the National Institute of Mental Health (NIMH) R01MH120227 (Perlis) and R01MH131687 (Sheridan and Perlis). The next generation sequencing was performed in Tufts University Core Facility Genomic Core, which received funding support from NIH (S10 OD 032203) for its Illumina NovaSeq sequencer.

## Competing Interests

Dr. Perlis has received personal fees from Circular Genomics, Genomind, Alkermes, and Atella for service as a scientific advisor, unrelated to the work described. He is a paid editor at JAMA+ AI and AI editor at JAMA Network-Open. The other authors have declared no competing financial interests in relation to the work described.

## Data Availability

Gene expression data will be made available for download from the NCBI Gene Expression Omnibus (https://www.ncbi.nlm.nih.gov/geo) upon publication. Additional data that support the findings of this study will be available from the corresponding authors upon reasonable request.

## Materials & Correspondence

Correspondence and material requests should be addressed to R.H.P or S.D.S.

## Notes

### Competing Interest Statement

The authors have declared no competing interest.

