## Supplemental Materials and Methods for "Functional Genomic Profiling of Schizophrenia-Associated Genes Reveals Key Microglial Regulators"

#### Description of Additional Supplemental Data Tables

**Supplemental Table 1.** Curated postmortem genes and prioritization of 30 pilot genes.

**Supplemental Table 2.** gRNAs used for primary screen/DRUG-seq and secondary screen.

**Supplemental Table 3.** Primary screen measurements – Phagocytic Index, eccentricity, solidity, cell# per well.

**Supplemental Table 4.** Secondary screen measurements – Phagocytic Index, eccentricity, solidity, IBA1 intensity, CD68 intensity, cell# per well.

**Supplemental Table 5.** List of DRUG-seq sequencing barcodes.

**Supplemental Table 6.** Differentially Expressed Genes (DEGs padj<0.1) by targeted gene.

**Supplemental Table 7.** Enriched Gene Ontology Biological Processes (GO-BP) per targeted gene.

**Supplemental Figures:**

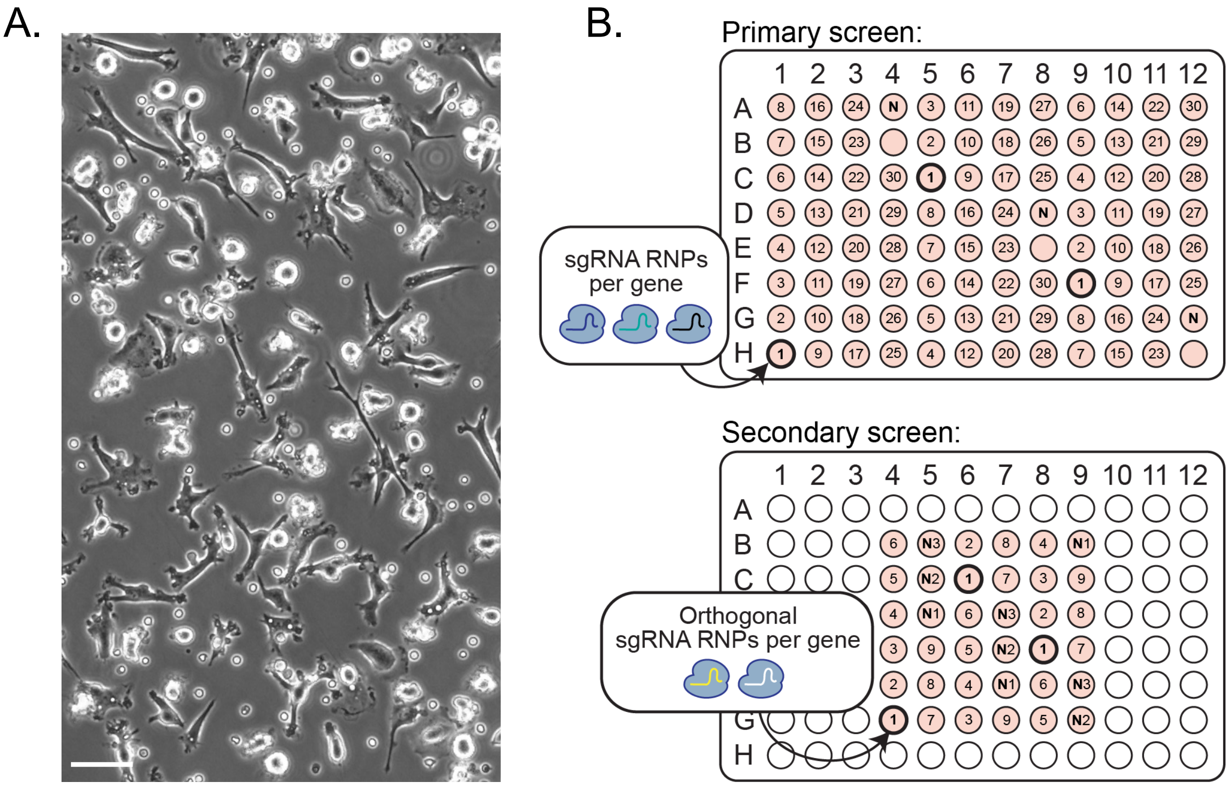


Figure S1. Functional genomics with PBMC microglia-like cells (piMGLCs). (a) Representative phase-contrast image of piMGLCs. Scale bar, 100 µm. (b) Image-based 96-well arrayed CRISPR functional synaptosome phagocytosis screening plate design. N indicates negative control non-targeting gRNAs.

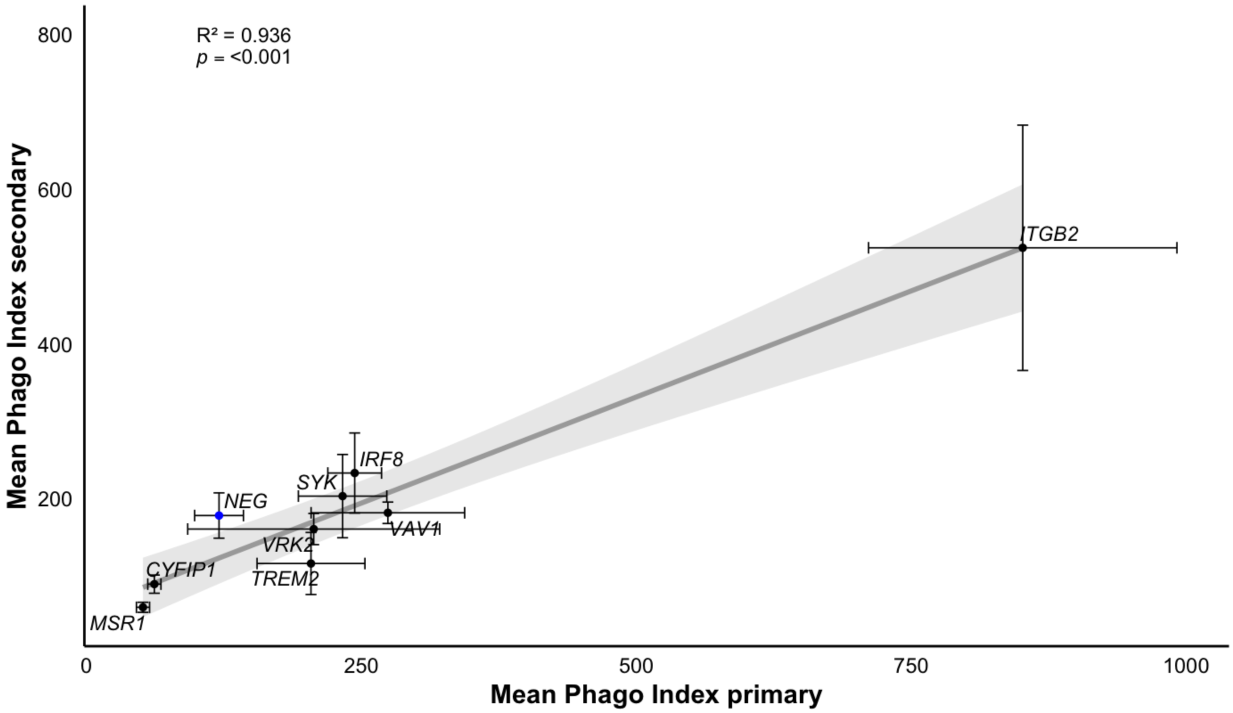


Figure S2. Correlation of phagocytic indices between primary and secondary screens excluding *ITGAM*. (*n*=3 replicate well mean cellular values, # cells per well in Tables S3 and S4). Error bars indicate SEM. Grey area indicates 95% confidence level.

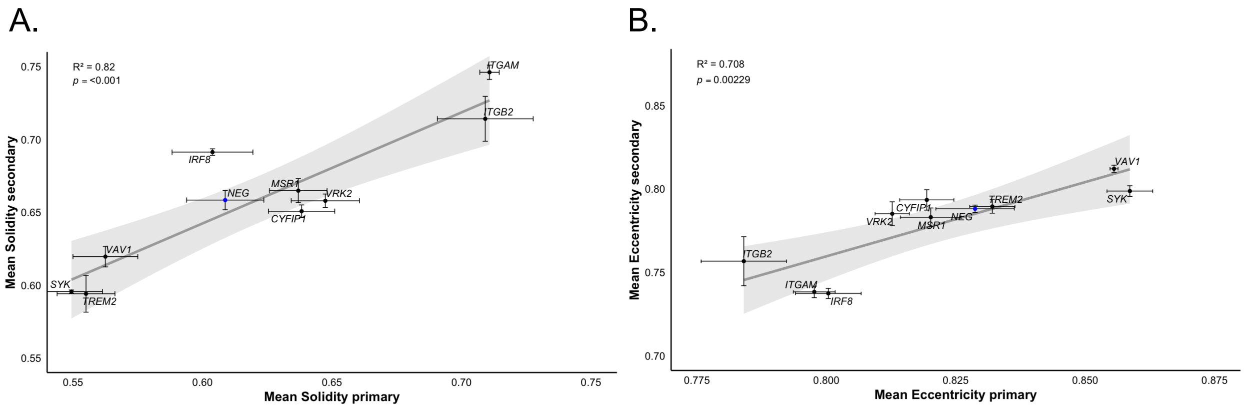


Figure S3. Morphology correlations between primary and secondary screens - secondary phenotypic confirmation screening confirms genes that modulate piMGLC morphology. Correlation of morphometric parameters (a) solidity and (b) eccentricity between primary and secondary screens for 9 CRISPR targeted genes selected from primary screen. (*n*=3 replicate well mean cellular values, # cells per well in Tables S3 and S4). Error bars indicate SEM, grey area indicates 95% confidence.

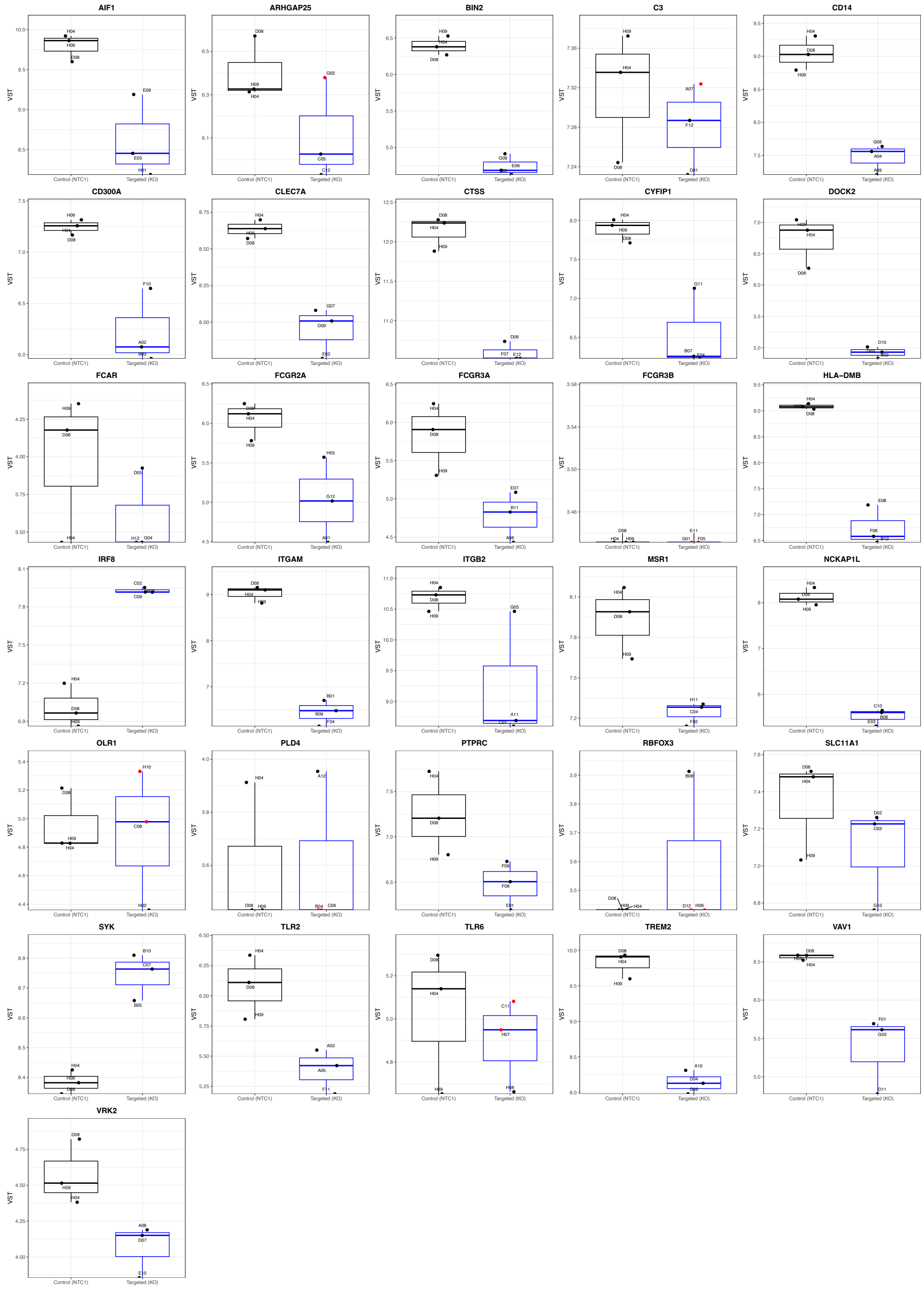

Figure S4.Variance Stabilizing Transform Normalized DrugSeq Counts for all target genes.

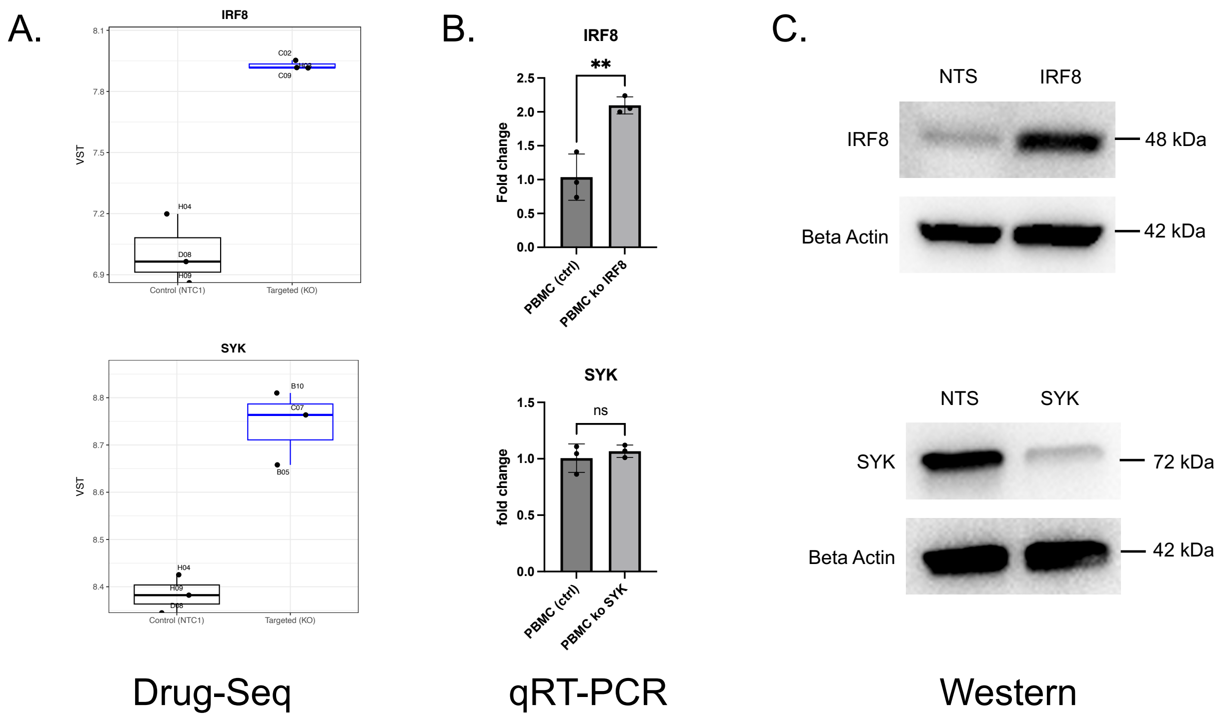


Figure S5. Differential expression of target genes *IRF8* and *SYK* as determined by (a) Variance Stabilizing Transformed Normalized DrugSeq Counts and (b) qRT-PCR. Protein expression of these target genes was determined by (c) Western blot analysis for negative control non-targeting sequence (NTS) or indicated gene gRNAs treated piMGLCs, respectively.

### Supplemental Methods:

#### Large-scale PBMC Isolation from Leukapheresis

Peripheral blood mononuclear cells (PBMCs) were isolated from a half leukapheresis pack (Leukopak, HemaCare/Charles River) as we have previously described.(1) Briefly, Leukopak contents were diluted 1:2 with HBSS (Thermo Fisher #14175095). 15ml of Lymphoprep Density Gradient Medium (Stemcell Technologies #07851) was added to 50ml conical tubes, and the diluted leukopak was layered on top. Tubes were centrifuged at 400g for 30 min (brake off). PBMC layers were collected, diluted to 45 ml with HBSS, and washed twice by centrifugation at 300g for 10 min (brake off), resuspending pellets in HBSS between washes. Cells were pooled, counted (Countess II, Thermo Fisher), and resuspended in Cryostor CS10 (Sigma-Aldrich #C2874) at 25 million cells/ml. After a final spin (300g, 10 min), pellets were resuspended in Cryostor CS10, aliquoted, and frozen in a Mr. Frosty container (Thermo Fisher #5100-0001) at -80°C before transfer to liquid nitrogen. Cells remained in Cryostor at room temperature for no more than 10 min prior to freezing; if longer processing was needed, suspensions were split and processed in batches.

#### Large-Scale Synaptosome Isolation from iPSC-Derived Neural Cultures

Human iPSCs were reprogrammed from fibroblasts and differentiated into neural progenitor cells (NPCs) as previously described for large scale neuronal culture differentiation and synaptosome isolation.(1-5) NPCs were expanded in neural expansion media (50% Advanced DMEM/F12, 50% Neurobasal, and neural induction supplement; Gibco: 21103-049, 12634-010, A16477-01), then seeded at 50 million cells per T1000 flask (Millipore, #PFHYS1008) for neuronal differentiation. Neuronal differentiation was performed in Neurobasal medium (Gibco #21103049) with N2 supplement (Stemcell Technologies SCT #7156), B27 supplement without Vitamin A (Gibco #12587010), NEAA (Gibco #11140050), penicillin/streptomycin, 1 μM ascorbic acid, 10 ng/mL BDNF and GDNF (Peprotech), and 1 μg/mL mouse laminin (Sigma #L2020), with weekly media changes for 8 weeks. Twenty-four hours before synaptosome isolation, cultures were switched to human astrocyte-conditioned medium (ScienceCell, #1811).

For synaptosome isolation, media was replaced with 1X gradient buffer (0.32M sucrose, 0.75mM $NaHCO_3$, 5mM Tris in Milli-Q water) plus HALT protease/phosphatase inhibitor (Thermo Scientific, #78440). Cultures were detached, homogenized (Bellco Glass, #1984-10015; 12 strokes/10ml), and centrifuged at 700g, 4°C, 10 min. Supernatants were saved and the pellets were resuspended in 1X gradient buffer to repeat the previous step once more. Supernatants were combined and transferred to high-speed centrifuge tubes (Beckman Coulter, #344058) to spin at 15000g, 4°C, for 15 minutes. Pellets were resuspended in gradient buffer and layered atop sucrose gradients (0.85M over 1.28M) in ultracentrifuge tubes. Gradients were spun at 26,500 rpm, 4°C, 2 hr (Beckman Coulter Optima L-90K, SW 32 Ti rotor). The synaptosome fraction (between 1.28M and 0.85M) was collected using a 5ml syringe with 16G needle, diluted 1:5 with 1mM $NaHCO_3$, and centrifuged at 20,000g, 4°C, 20 min. Final pellets were resuspended in gradient buffer plus 1 mg/ml BSA and aliquoted into cryovials (100ml/tube). Protein concentration was measured using the Pierce BCA assay (Thermo Scientific, #23225).

**Immunofluorescence**

Cells were fixed in 4% paraformaldehyde (Electron Microscopy Sciences, #15713S) for 15 min at room temperature. In 96-well plates, cells were washed three times with 100 μl Wash Buffer (PBS + 0.5% FBS). Cells were then incubated in 100 μl Block/Permeabilization Buffer (PBS + 0.5%

FBS + 0.3% Triton-X) for 1 hr at room temperature, followed by three washes with Wash Buffer. Primary antibodies (IBA1; Synaptic Systems #234009, 1:1000 and CD68; Abcam # ab213363, 1:100) diluted in Antibody Buffer (PBS + 0.5% FBS + 0.1% Triton-X) were added and incubated for 1 hr at room temperature or overnight at 4°C. Wells were washed three times, then incubated with secondary antibodies diluted in Antibody Buffer for 1 hr at 4°C. After three final washes, wells were imaged at 20X (37 image fields per well) using an IN Cell Analyzer 6000 (Cytiva).

**High Content Phagocytosis and Morphometric Image Analysis**

Confocal images were analyzed with CellProfiler (Version 4.2.1) (6) using the pipeline included in the github repository (https://github.com/mgb-cqh/crispr30). To summarize, images (37 images fields per well) were corrected for background signal, objects such as nuclei, cytoplasm, and synaptosomes were segmented, and features such as size, shape, and intensity were quantified and exported. Data were cleaned in R (R version 4.2.0, http://www.r-project.org), applying filters to exclude images with too few or too many cells, or images including over-segmentation. Phagocytic index was calculated as the sum of area of engulfed synaptosomes divided by number of cells per well.

Cell morphology was quantified using eccentricity and solidity metrics in CellProfiler, as previously described.(1, 6) Eccentricity was defined as the ratio of the distance between the foci of an ellipse fitted to each cell and the length of its major axis (range: 0–1), with higher values indicating elongated cells. Solidity was calculated for each cell as the ratio of the cell area to its convex hull area, with values near 1 signifying circular cells and lower values indicating branched or irregular morphologies.

**Multiplexed library preparation and RNA sequencing**

96-well multiplexed libraries (DRUG-seq (7)) were prepared using the MERCURIUS™ DRUG-seq kit (Alithea Genomics, 10841) as per manufacturer instructions. Briefly, cells were washed with 100 µL Dulbecco's PBS (Gibco, 14190-144) per well, lysed at 4°C with 20 µL/well 1x Cell Lysis Buffer, and clarified by centrifugation (300 × g, 5 min, 4°C). Up to 20 µL supernatant was transferred to a new 96-well PCR plate and stored at -80°C.

For reverse transcription, lysates were thawed and 10 µL was added to wells containing dry barcoded oligo-dT primers (Alithea Genomics, 10513). After resuspension, 10 µL RT Master Mix was added and reverse transcription performed (30 min 50°C, 10 min 85°C, hold at 4°C). First-strand cDNA was pooled, mixed with 7x DNA binding buffer (Zymo, D4003-1-L), and purified via Zymo-Spin IC column (C1004-50), eluting in 20 µL nuclease-free water. Excess oligo-dT was digested from 17 µL eluate using Exonuclease I, incubating 30 min at 37°C, 20 min at 80°C, and cooling to 4°C. Second-strand synthesis was performed immediately after (addition of 7 µL Second Strand Synthesis reaction mix, 20 min at 37°C, 30 min at 65°C, hold at 4°C).

Second-strand cDNA was purified with CleanNGS beads (Bulldog Bio, CNGS005; 0.6x ratio) using a PCR Strip Magnetic Separator (Permagen, MSR812). cDNA concentration was measured using Qubit 1X dsDNA HS Assay Kit (Invitrogen, Q33231); 50–60 ng cDNA was tagmented in a 20 µL reaction (with 4µL of provided Tagmentation Enzyme Buffer) for 7 min at 55°C, then purified with 0.6x CleanNGS beads. The resulting library was pre-amplified with provided kit UDI adapters (10 cycles: 98°C 10 s, 63°C 30 s, 72°C 1 min), then purified twice with 0.7x CleanNGS beads and eluted in 20 µL.

Final library quality and concentration were measured on an Agilent 5300 Fragment Analyzer with the HS NGS Fragment Kit (DNF-474-0500). Libraries were sequenced on a NovaSeq X Plus 1.5B flow cell (Read 1: 28 cycles, i7/i5: 8 cycles each, Read 2: 90 cycles).

**RNA-seq data analysis**

Raw paired-end FASTQ files were assessed with *FastQC* (v0.11.8) and adapters/low-quality bases removed using Trim Galore (v0.6.10). Post-trim quality was re-evaluated with *FastQC*. Trimmed reads were aligned to human reference genome *hg38* using *STAR* v. 2.7.11b (8) using "solo" mode. Unique molecular identifier count matrices were generated in this step by supplying the list of sequencing barcodes (Supplemental Data Table S5) to option *–soloCBwhitelist* with UMI de-duplication as described in the MERCURIUS DRUG-Seq kit documentation (Alithea Genomics, 10841). The resulting demultiplexed count matrix contained on average 1,514,089 ± 391,459 reads per sample and was used for downstream analysis.

Differential expression (DE) was performed with *DESeq2*.(9) Samples from each knockout (KO) gene well were contrasted to negative control (NTCs) wells. Genes with average copies per million < 2 in control wells were removed from analysis prior to model fitting. Models were fit with DESeq2, and log2 fold-changes were shrinkage-estimated with *apeglm*. Two different statistical criteria were applied. For the CRISPR-targeted gene itself**,** significance was determined using the nominal (unadjusted) *p*-value < 0.10 and raw (unshrunken) log2 fold-change values (Fig. 4**)**, under the rationale that expression of the perturbed gene is expected to change, reducing the likelihood of false positives relative to transcriptome-wide tests. For all other genes within each comparison, significance was assessed using FDR (Benjamini–Hochberg) adjusted padj < 0.10 and shrunken log2 fold-changes, appropriate for testing a large number of genes transcriptome-wide. Variancestabilized counts were used for visualization (Fig. S4). Functional enrichment of differentially expressed genes (DEG) was conducted for all CRISPR-targeted genes with >10 DEG with *clusterProfiler enrichGO*, with redundancy reduction by simplify (cutoff=0.7, by="p.adjust") where indicated. Original code developed for this study have been deposited in a public github repository (https://github.com/mgb-cqh/crispr30).

**RNA Extraction and qRT-PCR**

RNA was extracted from PBMC-derived microglia like cells (piMGLCs) lysed in Qiazol using the miRNeasy Micro kit (Qiagen, #217084), followed by on-column dsDNAse (Qiagen, #79254) treatment according to the manufacturer's instructions. Concentration was measured using the Qubit RNA High Sensitivity Assay kit (Invitrogen, # Q32852) on a Qubit 3.0 Fluorometer (Invitrogen, # Q33216). 38.64 ng of dsDNAse-treated RNA per sample was reverse transcribed using the Superscript III First-Strand Synthesis System (Invitrogen, #18080051) following the manufacturer's instructions. qPCR reaction was performed in triplicate using 2ng of cDNA. Each reaction contained 5µL of 2x Taqman Fast Advanced Master Mix (Invitrogen, #4444557), 4.5 µL of diluted cDNA and 0.25 µL of both FAM (IRF8 Hs00175238_m1; SYK Hs00895377_m1) and VIC (RPLP0 Hs00420895_gH) 20x Taqman Gene Expression Assay. Cycling was performed in the Quantstudio 7 Pro (Applied Biosystems, # A43183) with the following cycle conditions: 50°C for 2 min, 95°C for 2 min, 40x cycles of [95°C for 1 s, 60°C for 35 sec + single acquisition], 40°C for 10 s. Ct values above 35 were considered noise, and each gene of interest was normalized to the housekeeping control RPLP0 in the same well. The fold change was calculated using the 2^(-(deltadeltaCt)) method using the average of control samples as reference.

**Western blot Analysis**

For protein extraction, PBMC-derived microglia like cells (piMGLCs) were lysed with 1x RIPA buffer (Boston BioProducts, #BP-116TX) supplemented with protease inhibitor cocktail (Millipore Sigma, #1183617001), incubated on ice for 20 minutes, and centrifuged at 12,000 x g at 4 °C. The supernatant was collected, and protein concentration was determined using the Piece BCA Protein Assay (Thermo Scientific, #23225). Five micrograms (SYK) or 30 ug (IRF8) lysates were mixed with 1x Loading Dye (Boston BioProducts, #BP 111NR) and 355 mM β-mercaptoethanol (B-ME) and heated to 95 °C for 5 min. Precision Plus Protein Dual Color Standards (Bio-Rad, #1610374) were used for ladders. Samples were run on 10 % mini-PROTEAN gel (Bio-Rad, #456–1031) in 1x Tris-Glycine-SDS running buffer (Boston BioProducts, #BP-150) at 60 V for 30 minutes for stacking, followed by 85 V for 2 h. Protein was transferred to an Immun-Blot PVDF membrane (Bio-Rad, #1620177) in 1x transfer buffer (Boston BioProducts, #BP-190) using the BioRad Mini PROTEAN Tetra Cell at 125 V for 1 h. Membranes were blocked with 5 % non-fat dry milk (Millipore, #1.15363) in Tris-Buffered Saline with 0.1 % Tween-20 (Sigma, #P2287) (5 % milk TBST) for 1 h at room temperature. Primary antibodies (Rabbit anti-SYK, 1:5000, Cell Signaling, #13198T; Rabbit anti-IRF8, 1:1000, Cell Signaling, #5628T; Mouse anti-Beta Actin, 1:1000, Abcam, #ab8226) were diluted in 5 % milk TBST and incubated overnight at 4 °C. Membranes were washed with TBS and 0.1 % Tween-20. Secondary antibodies (Goat anti-Rabbit IgG (H+L) Secondary Antibody, HRP, 1:1000, Invitrogen, #31460; Goat anti-Mouse IgG (H+L) Secondary Antibody, HRP, 1:1000, Invitrogen, #62-6520) were diluted in 5 % milk TBST and incubated for 1 h at room temperature. Final washes were performed as described. Membranes were developed using SuperSignal West Atto Ultimate Sensitivity Substrate (Thermo Scientific, #A38554) and imaged using Bio-Rad ChemiDoc MP.
